# Parameterizing neural power spectra

**DOI:** 10.1101/299859

**Authors:** Matar Haller, Thomas Donoghue, Erik Peterson, Paroma Varma, Priyadarshini Sebastian, Richard Gao, Torben Noto, Robert T. Knight, Avgusta Shestyuk, Bradley Voytek

## Abstract

Electrophysiological signals across species and recording scales exhibit both periodic and aperiodic features. Periodic oscillations have been widely studied and linked to numerous physiological, cognitive, behavioral, and disease states, while the aperiodic “background” 1/f component of neural power spectra has received far less attention. Most analyses of oscillations are conducted on *a priori*, canonically-defined frequency bands without consideration of the underlying aperiodic structure, or verification that a periodic signal even exists in addition to the aperiodic signal. This is problematic, as recent evidence shows that the aperiodic signal is dynamic, changing with age, task demands, and cognitive state. It has also been linked to the relative excitation/inhibition of the underlying neuronal population. This means that standard analytic approaches easily conflate changes in the periodic and aperiodic signals with one another because the aperiodic parameters—along with oscillation center frequency, power, and bandwidth—are all dynamic in physiologically meaningful, but likely different, ways. In order to overcome the limitations of traditional narrowband analyses and to reduce the potentially deleterious effects of conflating these features, we introduce a novel algorithm for automatic parameterization of neural power spectral densities (PSDs) as a combination of the aperiodic signal and putative periodic oscillations. Notably, this algorithm requires no *a priori* specification of band limits and accounts for potentially-overlapping oscillations while minimizing the degree to which they are confounded with one another. This algorithm is amenable to large-scale data exploration and analysis, providing researchers with a tool to quickly and accurately parameterize neural power spectra.

## Introduction

In addition to being one of the first-ever observed features in human electrophysiology dating back to the original human electroencephalography (EEG) performed in 1929^1^, neural oscillations are widely-studied in neuroscience, with tens-of-thousands of publications to date. Close to a century of research has shown that oscillations may aid in coordinating interregional information transfer^2,3^, and suggest that oscillations influence a variety of cognitive, perceptual, and behavioral states^4,5^. Oscillatory dysfunctions have also been implicated in nearly every major neurological and psychiatric disorder^6,7^. Following historical traditions, the vast majority of the studies examining oscillations analyze canonical bands of interest, which are approximately defined as: delta (1-4 Hz), theta (4-8 Hz), alpha (8-12 Hz), beta (12-30 Hz), low gamma (30-60 Hz), high gamma (60-250 Hz), and fast ripples (200-400 Hz). Although all of these bands are frequently described as oscillations, standard approaches to analyzing these frequency bands fail to assess whether an oscillation—meaning rhythmic activity at a particular frequency—is truly present.

Given that there is a great deal of variability in oscillation center frequency across species^8^, age^9,10^, and cognitive/behavioral states^11-13^, it is easy to conflate power changes with center frequency differences (**Fig. 1**). This variability in oscillation characteristics is, at best, ignored by most approaches examining predefined bands and, at worst, can lead to misinterpretations of obtained results. For example, what is thought to be a difference in oscillatory band power could, in fact, reflect center frequency differences between groups of interest^14^ (**Fig. 1C**). Additionally, interpretation of band-limited power differences is confounded by the fact that oscillations are embedded within an aperiodic, 1/f signal that is also dynamic and may represent both background neural noise as well as physiologically relevant signals (**Fig. 1**). This means that power within a predefined frequency range does not necessarily imply oscillatory power. Because of this, a change in aperiodic slope between two groups—which might reflect tonic differences in excitation/inhibition balance^15^—or an event-related change in slope—such as seen in visual cortex^16^—might manifest as a simultaneous low-frequency power decrease and high-frequency increase, or vice versa (**Fig. 1C**). In this framing, changes in power ratios between bands may in fact reflect aperiodic slope differences totally free of any change in true oscillatory power in any band.

**Figure 1.**
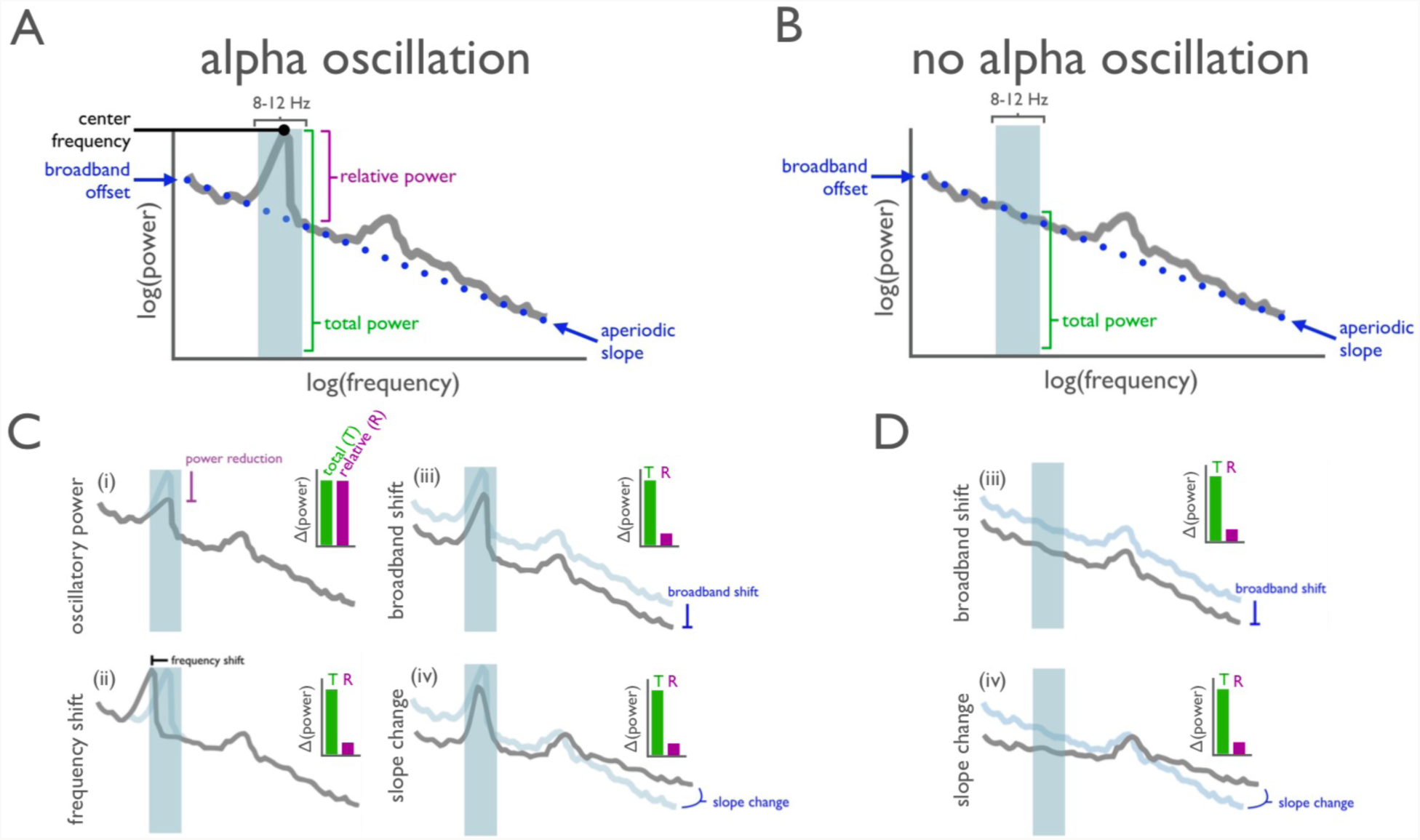
Overlapping nature of periodic and aperiodic spectral features. (**A**) Oscillations manifest as narrowband peaks in power above the aperiodic signal (blue dotted line)^25,28^, such as here, with the strong 8-12 Hz alpha peak (blue shaded region) and secondary beta peak (no marked). Narrowband filtering (*e.g.*, 8-12 Hz), without careful parameterization of the data, will give a numerical result even when, (**B**), there is no detectable oscillation present (here, artificially removed from (**A**)). (**C, D**) This can lead to misrepresentation and misinterpretation of physiological phenomena, because apparent changes in narrowband power can mimic several different physiological processes. These include: (i) a reduction in true oscillatory power^22,23^; (ii) a shift in the frequency of the oscillation^8-13^; (iii) a reduction in broadband power^17-19^; or; (iv) a change in the aperiodic slope^15,16,20,24-26^. In each case, *total* measured narrowband power is similarly changed (green bar), while only in the true power reduction case (i), is the 8-12 Hz oscillatory power *relative to* the aperiodic signal actually reduced (purple bar). Although the case shown in (i) is frequently assumed when narrowband power changes are observed, each of the alternative cases can also manifest as apparent oscillatory power changes, even when there is no oscillation present, such as in (**D**). Adjudicating between each of the physiological cases in i-iv, and between true and illusory oscillations, requires careful parameterization of the power spectrum.

In addition to the slope of the aperiodic signal, the offset of the aperiodic signal also likely carries physiological information, such as total population spiking of the neuronal population^17,18^. Fluctuations in the broadband offset may be linked to the fMRI BOLD signal^19^, making it a potentially crucial bridge between microscale and macroscale neurophysiological and cognitive features. Parameters of the aperiodic signal are dependent on cognitive and perceptual^16^ states, and are altered in aging^20^ and disease^21^. The fact that the aperiodic signal is itself dynamic and may index physiological features that are, at least partially, independent from the physiological generators of oscillations, strongly suggests that oscillatory power should be explicitly measured separately from this background.

To summarize, each of these four features—oscillation frequency^8-13^, oscillation power^22,23^, aperiodic broadband offset^17-19^, and aperiodic slope^15,16,20,24-26^—can and do change in behaviorally and physiologically meaningful ways, and may be inter-related^27^. Therefore, it is imperative that they are each carefully parametrized to avoid conflating them with one another, and to avoid confusing the physiological basis of “oscillatory” activity that may not be oscillatory at all. That is, changes in any and all of those four features can give rise to exactly the same change in total narrowband power, while the power of the periodic oscillation against the aperiodic signal need not necessarily change (**Fig. 1C,D**). For example, if activity is analyzed in a narrow band without considering the aperiodic signal, an apparent change in *e.g.*, 10 Hz alpha power may actually be due to a "see-saw rotation" of the overall PSD, with a pivot point at around 20-30 Hz (**Fig 1C.iv,D.iv**). In this scenario—which is has been observed to occur in a task-related manner in human cortex^16^—power in all frequencies <20 Hz will have decreased while power at frequencies >20 Hz will have increased. However, it would be a mischaracterization to say that there was a task-related decrease in the alpha band because that is not the signal feature that was truly altered.

Furthermore, reliance on *a priori* frequency bands may result in the inclusion of aperiodic activity from outside the true physiological oscillatory band—whose center frequency and bandwidth does not fall exactly within the *a priori* band—thus masking crucial behaviorally and physiologically relevant information (**Fig. 1C.ii**). Additionally, *a priori* filtering can give rise to changes in apparent oscillatory power that might not arise from a change in the oscillation, *per se*, but rather is caused by a shift in the aperiodic offset or slope (**Fig. 2A**). This can manifest as illusory oscillations where no oscillation exists (**Fig. 2B**). This is critical because many aperiodic signals—such as white noise, pink (1/f) noise, or even a single impulse function—have power at all frequencies despite there being, by definition, no periodic oscillation in the signal (**Fig. 2B**).

**Figure 2.**
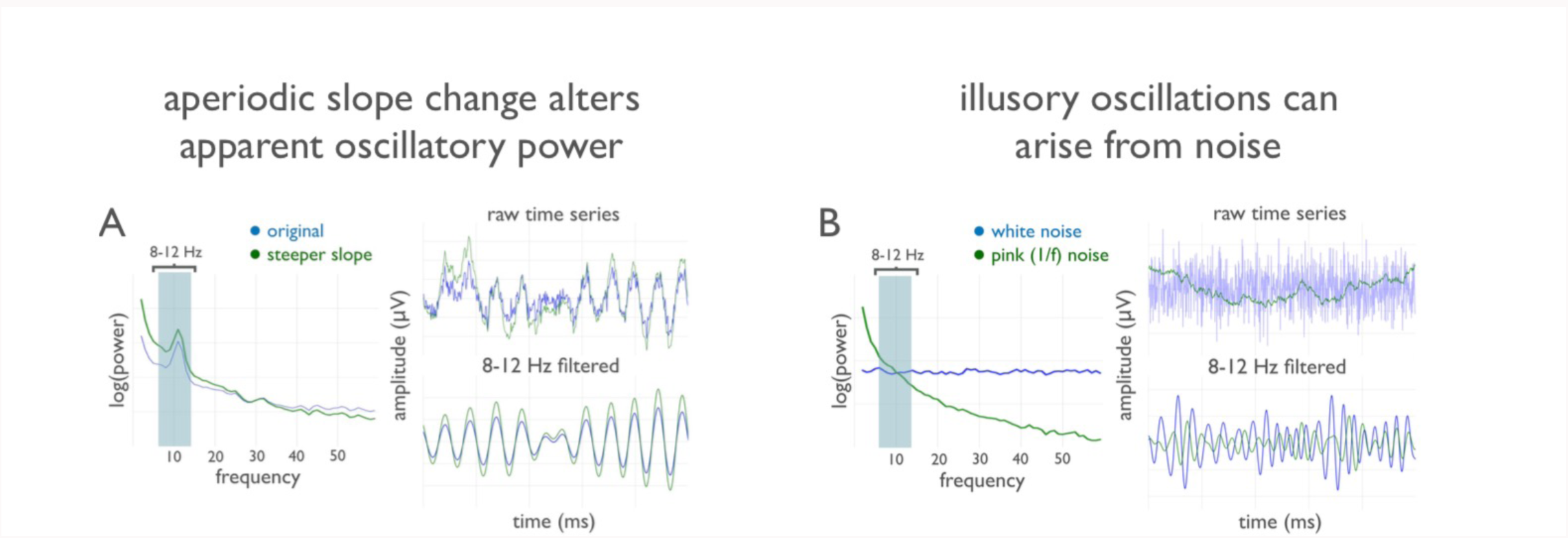
False oscillatory power changes and illusory oscillations. (**A**) A spectral change, such as seen in aging^20^—and artificially introduced in real data here—manifests as a dramatic difference between the time domain signals. This affects apparent narrowband power when an *a priori* filter is applied. This is despite the fact that the true oscillatory power *relative* to the aperiodic signal is unaffected. (**B**) Even when no oscillation is present, such as the case with the white and pink (1/f) noise examples here (blue and green, respectively), narrowband filtering gives rise to illusory oscillations where no periodic feature exists.

To overcome these limitations of narrowband analyses, we introduce an efficient algorithm for automatically parameterizing neural power spectral densities (PSDs) into periodic and aperiodic components. This algorithm extracts putative, periodic oscillatory components parameterized by their center frequency, power, and bandwidth, as measured from Gaussian model fits; it also extracts the offset and slope parameters of the aperiodic signal (**Fig. 3**). Importantly this algorithm requires no specification of narrowband regions to look for oscillations; rather the algorithm finds them automatically. While methods for identifying individual differences in oscillations exist, they are mostly restricted to identifying the frequency at which the power spectrum peaks within a specific sub-band^11^. This has resulted in a broad literature that has examined at least some aspects of variation within canonical oscillation bands, in particular the peak frequency within and across individuals, most commonly of the alpha band^11,13^. However, none of these methods address limitations regarding the use of canonical bands and the conflation of periodic activity with the periodic signal; additionally, they often assume only one peak within a particular band. Importantly, all of these methods measure *total* band power rather than power of the periodic oscillation *relative to* the aperiodic signal, further conflating the aperiodic and periodic processes.

**Figure 3.**
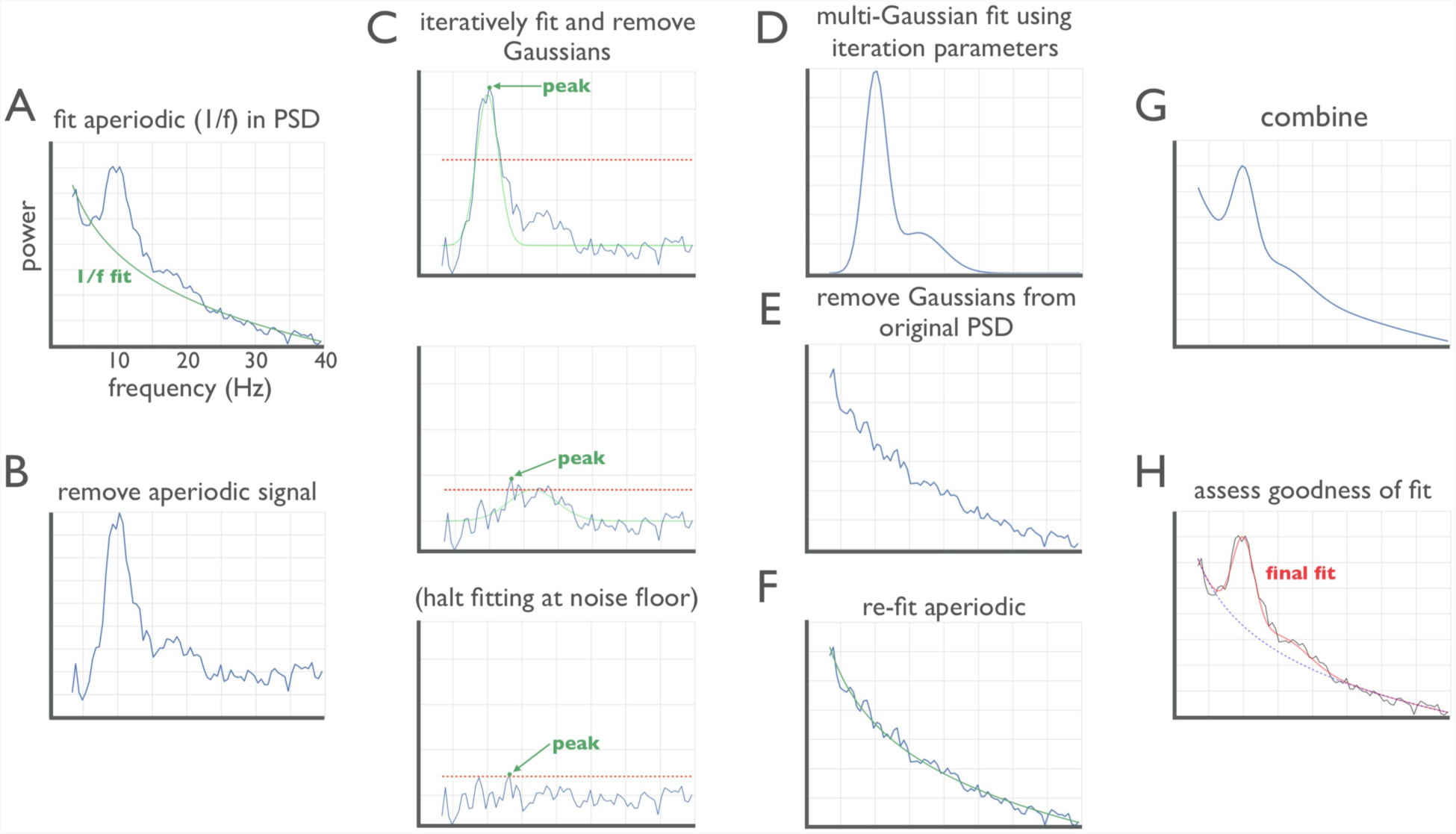
Algorithm schematic on real data. (**A**) The power spectral density (PSD) is first fit with an estimated aperiodic signal (green), defined by two parameters in *semilog-power* space: a slope and an offset. (**B**) The estimated aperiodic portion of the signal is subtracted from the raw PSD, the residuals of which are assumed to be a mix of periodic oscillatory peaks and noise. (**C**) The maximum (peak) of the residuals is found. If this peak is above the noise floor (2*std*; red dashed line) then a Gaussian (green) is fit around this peak based on the peak’s frequency, amplitude, and estimated bandwidth. The fitted Gaussian is then subtracted, and the process is iterated until the noise floor is reached (bottom). These values are used as seeds for the multi-Gaussian fitting in **D**. (**D**) Having identified the number of putative oscillations, based on the number of peaks above the noise floor, multi-Gaussian fitting is then performed on the aperiodic-adjusted signal from **B** to account for the joint power contributed by all the putative oscillations, together. (**E**) This multi-Gaussian model is then subtracted from the original PSD from **A**. (**F**) This is done to give a better estimate of the aperiodic signal—one that is less corrupted by the large oscillations present in the original PSD. (**G**) This re-fit aperiodic signal is combined with the multi-Gaussian model to give the final fit. (**H**) The final fit—here parameterized as a line (aperiodic signal) and two Gaussians (putative oscillations)— captures >99% of the variance of the original PSD. In this example, the extracted parameters for the aperiodic signal are: broadband offset = −21.4 au; slope = −1.12 au/Hz. Two Gaussians were found, with the parameters: (1) frequency = 10.0 Hz amplitude = 0.69 au, bandwidth = 3.18 Hz; (2) frequency = 16.3 Hz, amplitude = 0.14 au, bandwidth = 7.03 Hz.

## METHODS

Our parameterization method quantifies frequency characteristics of electro- or magnetophysiology data. This algorithm decomposes the original PSD into the aperiodic component and oscillatory peaks superimposed thereupon. While many methods can be used to calculate the PSD to be submitted to the algorithm for parametrization, here, for illustrative purposes, we use Welch’s method. The algorithm considers the PSD as the linear sum of an aperiodic signal and oscillations, or frequency regions of power over and above this aperiodic process, referred to as “peaks”. These peaks are considered to be putative oscillations, and are individually modeled as Gaussian functions. Each Gaussian is taken to represent an oscillation, whereby the three parameters that define a Gaussian are used to define the oscillation (**Fig. 3**).

This formulation fits the power spectrum as:

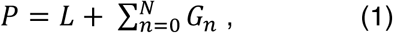

Where power, *P*, is a linear combination of the aperiodic signal, *L*, and there are *N* total ^Gaussians, *G*. Each *G*_*n*_ is a Gaussian fit to a peak, for *N* total peaks extracted from the power^ spectrum, modeled as:

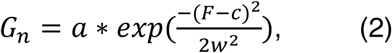

Where *a* is the amplitude, *c* is the center frequency, *w* is the bandwidth of the Gaussian, and *F* is the vector of input frequencies.

The aperiodic signal, *L*, is modeled using an exponential function in *semilog-power* space (linear frequencies and logged power values) as:

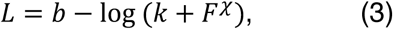

Where *b* is the broadband offset, χ is the slope, and *k* is the “knee” parameter, controlling for the bend in the aperiodic signal^29^, with *F* as the vector of input frequencies. Note that this formulation is equivalent to fitting a line in *log-log* space, when *k=0*, which we refer to as the “fixed” model. Fitting with *k* allows for modelling bends, or knees, in the aperiodic signal that are present in broad frequency ranges, especially in intracranial recordings^29^.

The final outputs of the algorithm are the parameters defining the best fit for the aperiodic signal and the *N* Gaussians. In addition to the Gaussian parameters, transformed parameters are also returned, in which we define: (1) center frequency as the mean of the Gaussian; (2) amplitude of the peak as the distance between the peak of the Gaussian and the aperiodic fit (this is different from the amplitude in the case of overlapping Gaussians), and; (3) bandwidth as 2*std*. Notably, this algorithm extracts all these parameters together in a manner that accounts for potentially overlapping oscillations; it also minimizes the degree to which they are confounded and requires no specification of canonical oscillation frequency bands.

To accomplish this, the algorithm first finds an initial fit of the aperiodic signal in *log*(power) by linear(frequency) space (**Fig. 3A**). This first fitting step is crucial, and not straightforward, as any normal fitting method such as linear regression, or even robust regression methods designed to account for the effect of outliers on linear fitting, can still be significantly pulled away from the aperiodic region due to the overwhelming effect of the putative oscillation peaks. To account for this, we introduce a procedure that attempts to fit the aperiodic aspects of the spectrum only. To do so, initial seed values for offset and slope are set to the amplitude of the first frequency in the PSD and −2.0 au/Hz, respectively (the latter being a good-enough guess for least-squares fitting based on empirical slopes^24^); these seed values are used to estimate a first-pass fit. The original PSD is then subtracted from this fit, creating a flattened spectrum, from which an amplitude threshold (set at the 2.5 percentile) is used to find the lowest amplitude points among the residuals, such that this excludes any regions with peaks that have high amplitude values in the flattened spectrum. This approach identifies only the data points along the frequency axis that are most likely to not be part of an oscillatory peak, thus isolating the parts of the spectrum that are most likely to represent the aperiodic signal (**Fig. 3A**). A second fit of the original PSD is then performed only on these frequency points, giving a better estimate of the aperiodic signal. This is, in effect, similar to approaches that have attempted to isolate the aperiodic signal from oscillations by fitting only to spectral frequencies outside of an *a priori* oscillation^20^, but does so in a more unbiased fashion. The percentile threshold value can be adjusted, if needed, but in practice rarely needs to be.

After the estimated aperiodic signal is isolated it is regressed out, leaving mostly the non-aperiodic features (putative oscillations) and noise (**Fig. 3B**). From this aperiodic-adjusted (*i.e.*, flattened) PSD, an iterative process searches for peaks that are then each individually fit with a Gaussian (**Fig. 3C**). Here, each iteration finds the highest peak in the aperiodic-adjusted (flattened) PSD. The location of this peak along the frequency axis is extracted, along with the peak amplitude. These stored values are used to fit a Gaussian around the peak, however a standard deviation is still needed. Thus, the standard deviation is estimated from the full-width, half-maximum (FWHM) around the peak by finding the distance between the half-maximum amplitudes on the left- and right flanks of the putative oscillation. In the case where there are two overlapping oscillations, this estimate can be very wide, so the FWHM is estimated as twice the shorter of the two sides. From FWHM, the standard deviation of the Gaussian can be estimated via the equivalence:

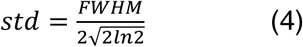

This estimated Gaussian is then subtracted from the flattened PSD, the next peak is found, and the process is repeated. By default, this oscillation-search step halts when it reaches the noise floor, based on a parameter defined in units of the standard deviation of the flattened spectrum (default = 2 *std*). Optionally, this step can also be controlled by setting an absolute amplitude, and/or a maximum number of Gaussians to fit. The amplitude thresholds (relative or absolute) determine the minimum amplitude beyond the noise floor that a peak must extend in order to be considered to be a putative oscillation. Once the iterative Gaussian fitting process halts, in order to handle edge cases, Gaussian parameters that heavily overlap (within 1.5 *std*), and/or are too close to the edge (<= 1.0 *std*) of the spectrum are then dropped. The remaining collected oscillation parameters for the *N* putative oscillations (center frequency, amplitude, and bandwidth) are used as seeds in a multi-Gaussian fitting method (Python: `scipy.optimize.curve_fit`). Each fitted Gaussian is constrained to be close to (within 1.5 *std*) of its original guessed Gaussian. This process attempts to minimize the square error between the flattened spectrum and *N* Gaussians simultaneously (**Fig. 3D**).

This multi-Gaussian model is then subtracted from the *original* PSD, in order to isolate an aperiodic signal from the modeled and parameterized oscillatory peaks (**Fig. 3E**). This oscillations-removed PSD is then re-fit, allowing for a more precise estimation of the aperiodic signal (**Fig. 3F**). This, when combined with the equation for the *N*-Gaussian model (**Fig. 3G**), gives a highly-accurate parameterization of the original PSD using few features (**Fig. 3H;** in this example, >99% of the variance in the original PSD is accounted for by the combined aperiodic + periodic model). Goodness-of-fit is estimated by comparing each fit to the original power spectrum in terms of the RMSE error of the fit as well as the *R^2^* of the fit. Optional parameters allow for tuning algorithmic performance based on different datasets, such as invasive LFP versus M/EEG. These optional parameters can define: (1) the maximum number of peaks; (2) limits on the possible bandwidth of extracted peaks, and; (3) absolute, rather than relative, amplitude thresholds.

Code for this algorithm is available as an open-source Python package, with support for Python >= 3.5, under the Apache-2.0 license, and is available on the Python Package Index^ii^. The package includes documentation and a test-suite, with a series of tutorials also available on the project GitHub page^iii^. Its package dependencies are limited to *numpy* and *scipy* (>= version 0.19). On contemporary hardware (3.5 GHz Intel i7 MacBook Pro), a single PSD is fit in approximately 10-20 msec. Because each PSD is fit independently, this package has support for running in parallel across PSDs to allow for high-throughput parameterization.

## DISCUSSION

Despite the ubiquity of oscillatory analyses—tens-of-thousands of peer reviewed publications indexed in PubMed—there are several analytic assumptions and potential artifacts that significantly impact the physiological interpretation of previous oscillatory research. In brief: 1) oscillations should be measured relative to the aperiodic (1/f) signal because, strictly speaking, oscillations are defined as any regions of the power spectrum that rise above the 1/f background^28^ (**Fig. 1**); yet this is rarely done, and; 2) most tools for quantifying oscillations assume that oscillations exist even though they may not even be present in the signal (**Fig. 2**); verification of the presence of an oscillation is also rarely done. Without careful parameterization *of all the components of the power spectrum*, it is easy to misinterpret the physiological relevance of any narrowband signal changes (**Fig. 1**).

To address this problem, we introduce a novel method for algorithmically extracting oscillatory components in electro-and magnetophysiological data. This algorithm addresses often-overlooked problems in cognitive and systems neuroscience. In particular, implicit reliance on canonical frequency bands can lead to both false positive, and false negative, results. For example, apparent group differences in oscillatory power may be the result of shifts in the center frequency of the oscillation (**Fig. 1C**) and not changes in true oscillation power. This can be illustrated by aging research wherein it is well-accepted that alpha frequency decreases with aging, yet there have also been reports of a decrease in alpha power with age^9,10^. If it is the case that younger adults have a 10 Hz alpha center frequency, while older adults have an equally high-power alpha that has slowed to 8 Hz, canonical frequency band analysis in the 8-12 Hz alpha range will give the false appearance of lower alpha power in older, relative to younger, adults due to the alpha oscillation moving outside the *a priori* alpha band, despite the fact that power need not have changed.

In another example, changes in the aperiodic signal, as seen with aging^20^ and behavior^16^, will shift total narrowband power despite the fact that power in a narrowband *oscillation* has not changed relative to the aperiodic process^7^ (see **Fig. 2A**). Note that the see-saw rotation phenomenon of the aperiodic signal parsimoniously explains the ubiquitous negative correlation between low frequency (<30 Hz) and high frequency (> 40 Hz) signals^30^. That is, when a full spectral parameterization is performed, rather than multiple narrowband analyses, it becomes clear that, rather than there being multiple interacting oscillatory processes such as an alpha oscillator and a gamma oscillator operating in a push/pull fashion, there may only be one physiological process that is changing: the slope of the aperiodic signal^7^. A change in the aperiodic slope also manifests in the time-domain as oscillatory amplitude and raw voltage differences (**Fig. 2A**). Given that such differences in the slope of the aperiodic signal have been observed across many different groups, such as in aging^20^ and disease^21^, it may be that differences in time-domain averaging based analyses, such as with event-related potentials or event-related spectral perturbations, may be partially explainable by aperiodic slope differences between groups or across conditions.

There are currently several algorithms for identifying oscillations in specific ways that have attempted to address some of these concerns individually, but never conjointly. In particular, an approach called BOSC (Better OSCillation Detector)^31^ is akin to the one presented here, beginning by fitting a linear regression to the *log-log* PSD to estimate the aperiodic signal. This is used to determine a power threshold, which is then used in combination with a duration threshold to define oscillations in wavelet-based decompositions of the time series data. However, a significant limitation of this and other similar methods is that a simple linear fit of the background spectrum can be significantly skewed by the presence of oscillations—especially large oscillations—and therefore mischaracterizes the aperiodic signal. This suboptimal aperiodic signal fit, in turn, hampers oscillation detection because the background fit is used to set a power threshold for extracting oscillations. Another similar approach is the irregular-resampling auto-spectral analysis (IRASA) method, which seeks to explicitly separate the periodic and aperiodic components through a resampling procedure^32^. Though conceptually similar, this resampling method is computationally much more expensive, prohibiting large-scale deployment, and has trouble separating large amplitude oscillations as it tends to blur them in frequency space. Other methods, such as lagged-coherence^33^, offer time-series analysis that is designed to differentiate rhythmic activity from transients, while principle component variants fail to separate periodic and aperiodic features, and require manual component selection^18^. While we show that the aperiodic signal is of significant physiological and behavioral interest, all of these methods treat it as a nuisance variable rather than a feature to be explicitly modeled. Further, none of these techniques offer full parameterization, making large-scale analysis and data aggregation difficult. Our algorithm addresses these limitations through improved background fitting, using an iterative approach that considers the background and oscillations together.

Traditional canonical frequency band analyses commit researchers to tacit acceptance of predefined oscillatory bands having a functional role, rather than considering the underlying physiological mechanisms that may generate different spectral features. Further, they fail to address inter-individual differences. For example, variations in peak-frequencies within oscillation bands have functional correlates and are of theoretical interest^13^. By providing a tool to more precisely parameterize such features, further study of such features is facilitated. In addition, recent advances in cross-frequency coupling analyses, such as phase-amplitude coupling (PAC), have provided a powerful means for probing the potential mechanisms of neural communication^3,34-36^. These analyses typically rely on canonical frequency bands, which is problematic given that true multiple-oscillator PAC is known to exhibit different phase coupling modes as a function of cortical region^35,37^. Additionally, the appearance of interacting oscillators can just as easily arise as an artifact of narrowband signal processing methods^38,39^. With proper parameterization of neural power spectra in such a way as to first characterize oscillatory components, it may be possible to identify phase coupling modes across brain regions, task, and time, thus increasing the specificity and accuracy of cross-frequency coupling analyses.

Relying on *a priori* frequency bands and averaging spectral features can also blur critical variability. There may exist a wide range of low gamma frequencies within subjects such that averaging across those bands decreases overall statistical power^40^. Furthermore, specific frequency bands, such as alpha, have been linked to a wide-array of sometimes-conflicting processes, such as inhibition or cortical potentiation^22^, periodic sampling^41^, and prediction^42^. It may be that many different physiological processes—including changes in the slope or offset of the aperiodic signal, and periodic oscillatory changes—are being conflated^43^, resulting in many different physiological processes being grouped together as “oscillatory alpha”^44^.

Our approach is a principled method for quantifying the neural power spectrum. Such an algorithm may allow us to better link macroscale electrophysiology to microscale synaptic and firing parameters^45-47^, providing a better understanding of the relationship between microscale synaptic dynamics, mesoscale LFP and ECoG/iEEG, and the macroscale EEG and MEG^48^. Our parameterization approach increases analytical power by disentangling the aperiodic and periodic components of neural power spectra. This allows researchers to take full advantage of the rich and meaningful variability present in neural field potential data, rather than treating that variability as noise to be averaged away. This approach has the potential to provide greater insight into both the physiological mechanisms underlying oscillations, as well as the role that oscillatory variability may play in explaining individual differences in cognitive functioning in health, aging, and disease. Finally, because of the speed and ease of the algorithm, and interpretability of the fitted parameters, this tool opens avenues for the high-throughput, large-scale analyses that will be critical for data-driven approaches to neuroscientific research^49^.

## Acknowledgements

We thank S.R. Cole, R. van der Meij, B. Postle, and T. Tran for invaluable discussion, usability testing, and comments. Haller is supported by a National Science Foundation (NSF) Graduate Research Fellowship Grant DGE1106400. Sebastian is supported by a UC San Diego Frontiers of Innovation Scholars Program fellowship. Gao is supported by the Natural Sciences and Engineering Research Council of Canada (NSERC PGS-D), UC San Diego Kavli Innovative Research Grant (IRG), Frontiers for Innovation Scholars Program fellowship, and a Katzin Prize. Knight is supported by a National Institute of Neurological Disorders and Stroke (NINDS) Grant R37NS21135. Shestyuk is supported by a National Institute of Mental Health (NIHM) Grant F32MH75317. Voytek is supported by the Whitehall Foundation, a Sloan Research Fellowship, and the National Science Foundation under grant BCS-1736028. The authors declare no competing financial interests.

## Footnotes

i https://github.com/voytekresearch/fooof/

ii https://pypi.python.org/pypi/fooof/

iii https://github.com/voytekresearch/fooof/

